# Early birds, late owls, and the ecological role of intra-population chronotype variation

**DOI:** 10.1101/2020.08.09.243261

**Authors:** Ella Royzrakh-Pasternak, Tamar Dayan, Ofir Levy, Noga Kronfeld-Schor

## Abstract

While the molecular mechanisms underlying variation in chronotypes within populations have been studied extensively, the ultimate selective forces governing it are poorly understood. The proximate cause is variation in clock genes and protein expression, which produces variation in tau (period length of the circadian clock), with early individuals having shorter tau. We studied within-population variation in foraging activity times of two *Acomys* species in the field. This variation manifested in a regular and consistent sequence of individual foraging activity that is positively and strongly correlated with variation in tau. Thus, variation in circadian clock period length (tau) appears to be the mechanism underlying the regular pattern of intraspecific temporal partitioning. Late chronotypes also spent more time torpid than earlier ones, suggesting an energetic cost to this strategy and possible tradeoffs. We suggest that variation in tau is an adaptive mechanism to reduce competition between individuals within a population.

## Introduction

The diel activity patterns of animals have attracted significant scientific interest focusing on their evolution, ecological significance, regulating mechanisms, and impacts of anthropogenic activities (1–5). The evolution of diel rhythms is driven by the 24 h light-dark cycle and its derivatives, including both environmental conditions and ecological interactions. In the past two decades a growing volume of research has addressed interspecific niche partitioning along the diel axis, with a multidisciplinary approach that involves community ecology, evolutionary biology, and biological rhythms research (2, 4). This research ranged from macroevolutionary patterns to studies of behavioral plasticity in response to human activity and artificial illumination and their bearing on species coexistence (e.g., (5, 6)).

Concurrently biological rhythms research has developed dramatically and while a significant part of it focuses on humans and model organisms, the past decade has seen a growing literature on animal biological rhythms. The 24 h light and dark cycle is a strong and predictable environmental pattern; daily rhythms in behavior and physiology are governed by a biological clock that is synchronized to this universal cue. When biological clocks synchronize or entrain to the environment (usually to the light/dark cycles), they not only show the same period as the zeitgeber (cue) cycle (24 h), but also establish a stable phase relationship with the zeitgeber, called the phase of entrainment (7, 8).

In spite of millions of years of natural selection to a strong universal cue, a significant variation in the period length of the internal biological clock occurs, resulting from variation in clock genes and protein expression (9, 10); consequently different individuals synchronize differently to the light/dark cycle, resulting in variability in their phase of entrainment relative to the environmental cue or zeitgeber (e.g., sunrise), called chronotypes. Chronotype describes the manifestation of daily rhythms, which are the outcome of an interaction between the endogenous circadian clock and environmental conditions (7, 11). They correlate with the free-running period of the circadian clock (the endogenous circadian clock period under constant environmental conditions, called tau), which has a significant genetic component, with early chronotypes displaying a shorter tau than late chronotypes (9, 12–14). Here we investigate the role that this genetic and phenotypic variation may have in the coexistence of individuals within a population and hence the selective forces that may be at play maintaining this variation.

Individual variation in daily rhythms under free-living conditions has been the focus of intensive research in humans during the last decades, with earlier individuals described as larks and late individuals as owls (7, 9, 13). Variability in chronotypes was also described in different animal species, including rodents, birds, fishes, insects and others, but these experiments were conducted almost exclusively under controlled laboratory conditions; very few studies attempted to address variation of chronotypes in natural populations of species other than humans. Even fewer studies (14–16) focused on the selective forces driving this variation which are yet to be understood (8). Several studies in recent years have demonstrated intraspecific patterns of temporal partitioning that implicated intraspecific competition (17, 18). We studied intraspecific variation in chronotypes in two coexisting wild rodent species, its adaptive significance and proximate underlying mechanisms, and the selective forces driving it.

In an earlier study we found regular intraspecific patterns of foraging activity among golden spiny mice (*Acomys russatus*), with specific individuals tending to arrive and forage early (early chronotype) and others to arrive and forage later (late chronotype) during the day (19). Persistence of intraspecific variation in chronotypes in natural populations suggests that a specific chronotype entails benefits or costs which depend on environmental conditions, and have fitness consequences. Under our field experimental setup, where food was supplemented and replenished every morning, we found that early individuals spent less time torpid (controlled reduction of body temperature (20, 21)) than late individuals (19). As use of torpor was related to a decrease in reproductive success in this species (22), it appears that early chronotypes had a selective advantage in this study system. Intra-population morphological variation has been studied empirically with a theoretical basis that related it to intra-specific competition, with variation related to resource breadth and availability (23). In the current study we hypothesized that in a similar manner, foraging order is determined by chronotype, which in turn is determined by tau, and that intra-population variation in tau, as a proximate mechanism driving variation in chronotype (and hence foraging sequence), will be likewise selected for to reduce resource use overlap and thus reduce intraspecific competition at the diel niche axis. To test this hypothesis, we measured the order of arrival to a foraging patch (foraging sequence) and use of torpor in golden and common spiny mice (*A. cahirinus*) under semi-natural conditions, and the free-running period lengths (tau) of the foraging individuals under controlled laboratory conditions.

## Results

The experiment yielded a total of 550,000 foraging logs and 300,000 body temperature records. For one common spiny mouse (from enclosure 3) there were no body temperature data, and three golden spiny mice (two from enclosure 3 and one from enclosure 2) were not recaptured and consequently we could not measure their torpor use or period length; these individuals were excluded from the data analysis.

### Common spiny mice

Individuals from the same enclosure entered the foraging patches in a constant order during the experimental period (Fig 1A). The free running period length (tau) of common spiny mice ranged between 23.7 h to 24.3 h (Fig. 2A).

**Figure 1:**
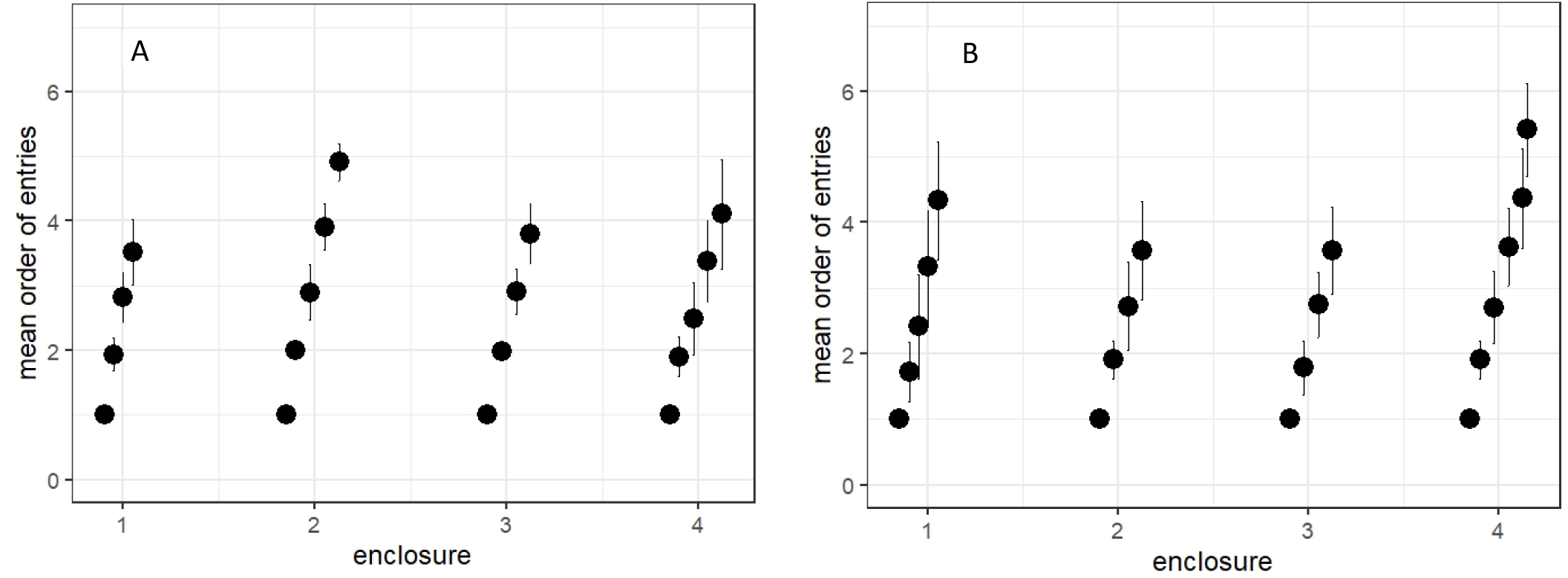
The mean order of entry (±SE) of common (A, n=18 individuals, 4 enclosures (replicates)) and golden (B, n=19 individuals, 4 enclosures (replicates)) spiny mice in each enclosure (1-4). Each point represents a different individual. One common spiny and three golden spiny mice were excluded from the data analysis, due to the reasons mentioned in the Results, p. 4). Source files of all the data used for the analysis are available in the Figures 1a, 2a, 3a first visit and tau *A. cahirinus* and Figures 1b, 2b, 3b first visit and tau *A. russatus* source data files.

**Figure 2:**
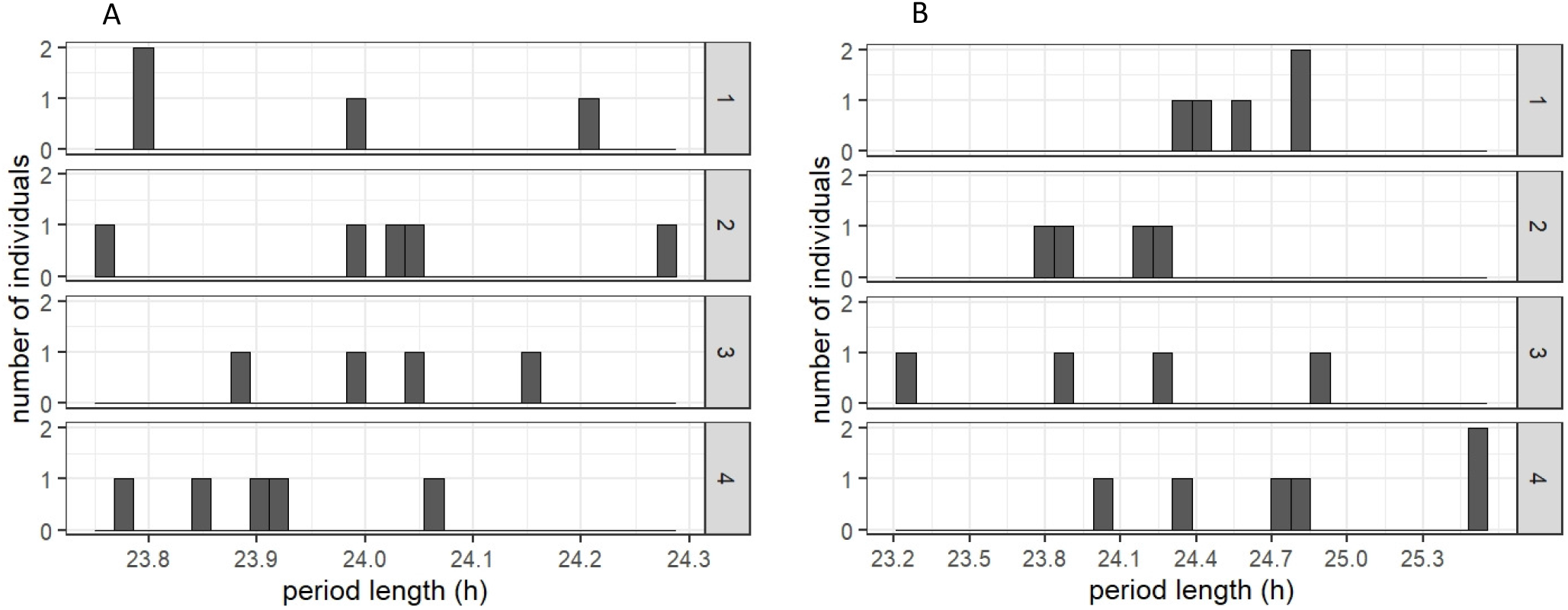
Distribution of free running period length (tau) in common (A, n=18 individuals, 4 enclosures (replicates)) and golden (B, n=1 individuals, 4 enclosures (replicates)) spiny mice individuals in each enclosure separately. One common spiny and three golden spiny mice were excluded from the data analysis, due to the reasons mentioned in the Results, p. 4). Source files of all the data used for the analysis are available in the Figures 1a, 2a, 3a first visit and tau *A. cahirinus* and Figures 1b, 2b, 3b first visit and tau *A. russatus* source data files.

Our statistical models suggested that individuals that had shorter period length (*τ*) arrived earlier to the foraging patches (Estimate= 0.6741, SD= 0.2353; t-value= 2.865; DF=16; p-value=0.00418, ΔAIC=4.2451, N=18; Fig.3A).

**Figure 3:**
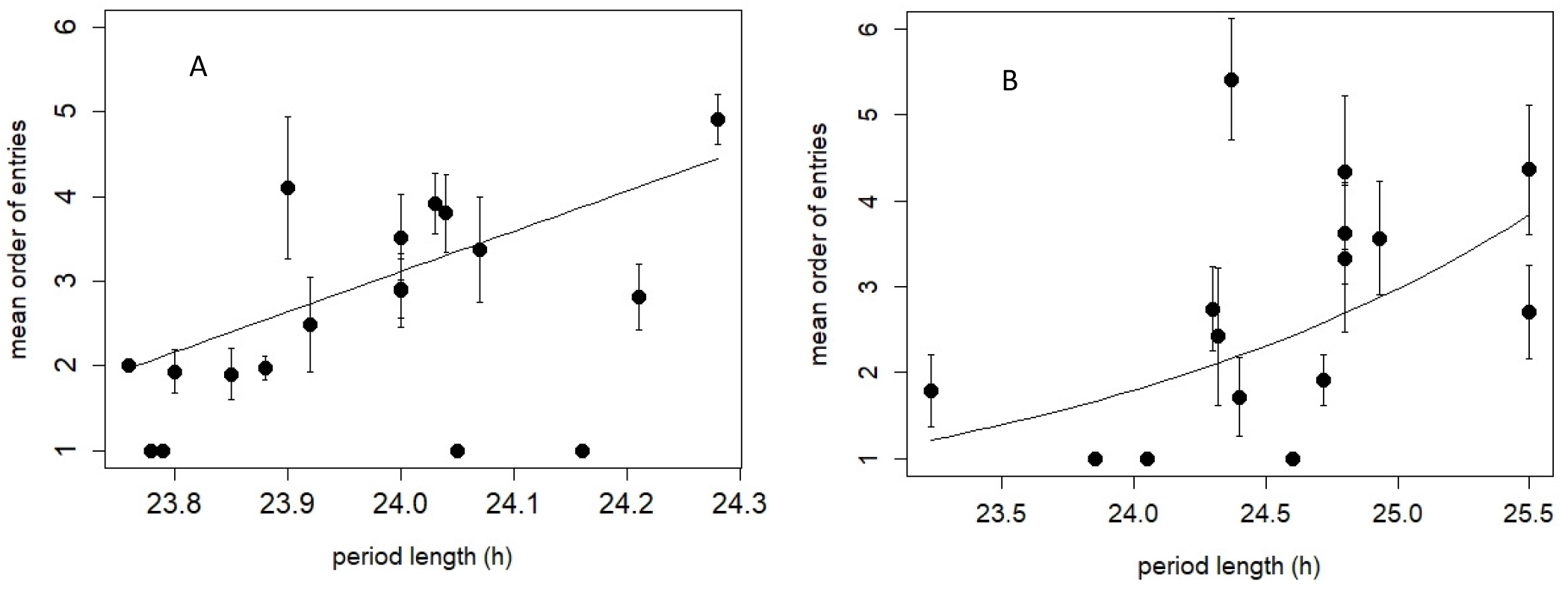
The correlation between individual mean daily order of entries (±SD) and period length (h) in common (A, n=18 individuals, 4 enclosures (replicates), p-value=0.00418, GLMM) and golden (B, n=15 individuals, 3 enclosures (replicates), p-value=0.0564, GLMM) spiny mice. The four golden spiny mice in enclosure # 2 were excluded from the data analysis due to the reasons mentioned in the Methods, p.5. Source files of all the data used for the analysis are available in the Figures 1a, 2a, 3a first visit and tau *A. cahirinus* and Figures 1b, 2b, 3b first visit and tau *A. russatus* source data files.

Most of the common spiny mouse individuals (82.35%) used torpor for fewer than 60 minutes during the day on average and (71.42% out of them used torpor for fewer than 20 minutes). The rest of the individuals (17.64%) used torpor for fewer than 80 minutes a day on average.

The period length did not affect the daily torpor usage of the common spiny mice (Estimate=−0.3880, SD= 0.2829; t-value= −1.372; p-value= 0.17; DF=15, ΔAIC=−0.16, N=18; Fig.4A), probably due the overall low torpor usage of the common spiny mice. In both tests, we used Generalized Linear Mixed Effect Models (GLMM), as indicated in the Methods section, page 12.

**Figure 4:**
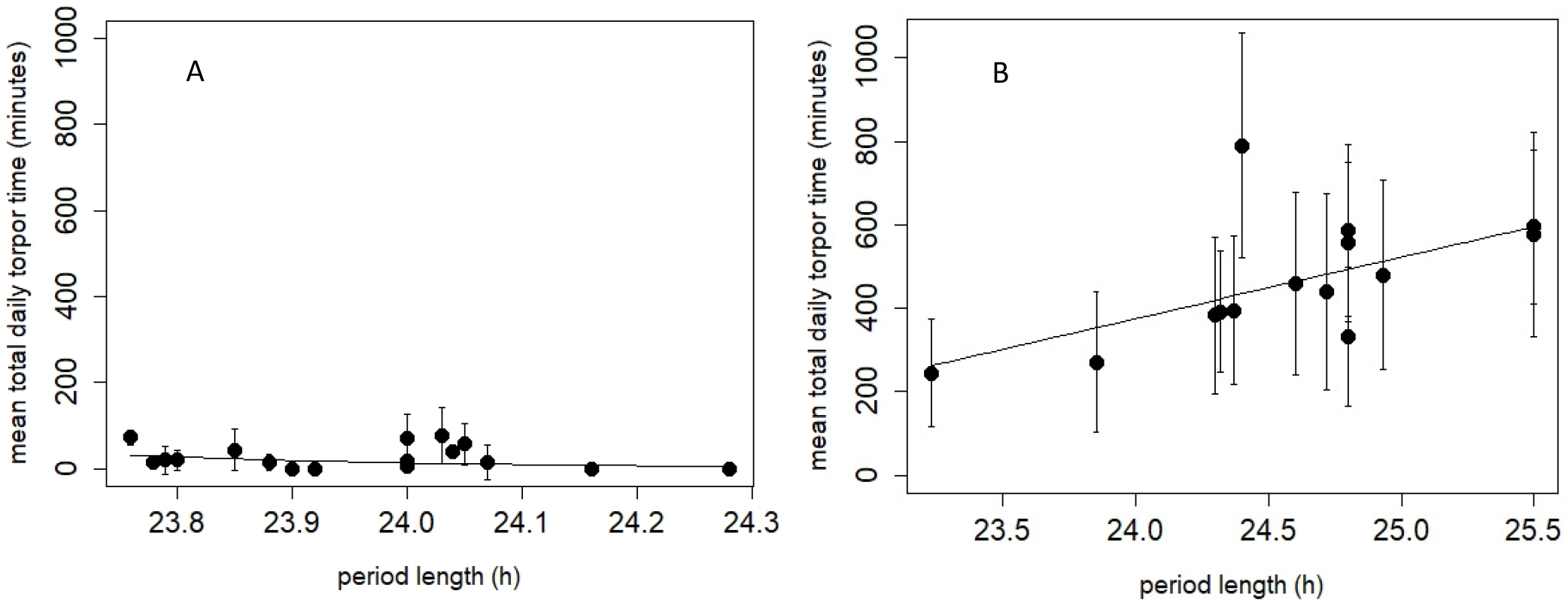
The correlation between the individual mean daily torpor time (minutes; ±SD) and the period length (h) in common (A, n=18 individuals, 4 enclosures (replicates), p-value=0.17, GLMM) and golden (B, n=15 individuals, 3 enclosures (replicates), p-value<0.001, GLMM) spiny mice. The four golden spiny mice in enclosure # 2 were excluded from the data analysis due to the reasons mentioned in the Methods, p.5. Source files of all the data used for the analysis are available in the Figures 4a first visit and torpor *A. cahirinus* and Figure 4b first visit and torpor *A. russatus* source data files.

### Golden spiny mice

Individuals from the same enclosure entered the foraging patches in a constant order during the experimental period (Fig 1B). Furthermore, the frequency distribution of the period length of the golden spiny mice individuals is dispersed between 23.2 h to 25.4 h (Fig. 2B). In enclosure # 2 where four golden spiny mice were placed, two pairs of individuals overlapped almost entirely in tau, and the four individuals were fairly close in tau as well (Fig 2B). Hence no correlations could be discovered and this enclosure was excluded from the following analyses.

Individuals that had shorter period length (*τ*) arrived earlier to the foraging patches (Estimate= 0.3036, SD= 0.1591; t-value= 1.908; p-value= 0.0564; DF=13; ΔAIC=1.1721, N=15; Fig.3B). The period length had a significant positive effect on the daily torpor use of golden spiny mice (Estimate= 86.55, SD= 19.91; t-value= 4.346; DF=12; p-value<0.001; ΔAIC=5.68, N=15; Fig.4B). The daily time spent torpid increased by 2.4 minutes with every change of 1 minute in the period length. In both tests, we used Generalized Linear Mixed Effect Models (GLMM), as indicated in the Methods section, page 12.

## Discussion

We found a constant order of arrival to the food patch in both *A. russatus* and *A. cahirinus*: individuals of both species tended to arrive to the food patch in the same order along the experimental period. In seven of the eight population in the enclosures, we also found that the order of arrival is positively correlated with the period of the internal biological clock (tau), measured under controlled conditions in the laboratory. In the *A. russatus* population in enclosure #2 we found no significant positive correlation, and hence removed it from further analysis. Finally, in *A. russatus* foraging sequence correlated with torpor use, an energy conserving mechanism. Research suggests that tau has a significant genetic component (9, 12–14), so it appears that the order of arrival to forage in a food patch, returns in energy acquisition, and the consequent energy conserving strategy required, are strongly influenced by the genetic makeup of the individual. We note that the entire pool of mice studied constitutes random sampling by trapping the local free-living spiny mouse populations, while the individuals populating specific enclosures are merely a randomly assigned sub-sample of this pool. The analysis of all studied individuals within species yielded clear correlations. However, as the numbers of individuals within species populating the enclosures are limited (4–6), they cannot be taken to fully represent tau in the native population. Even with this limitation, a clear pattern emerges in seven of eight populations. We thus suggest that in the field, intra-population variation in tau, as a proximate mechanism for driving variation in chronotype, is selected for to reduce resource use overlap or intraspecific aggression (see below) and thus to reduce intraspecific competition at the diel niche axis.

While use of time as a niche axis for reducing interspecific competition has received considerable attention during the last decade (reviewed elsewhere (3, 4)), it was seldom addressed in relation to intraspecific interactions. We have previously shown that activity during different parts of the diel cycle facilitates coexistence between the two spiny mouse species in this system (2, 24). Being active at different times exposes the two omnivorous species to different prey worlds and thus reduces resource use overlap (24, 25). It is possible that in the same manner, individuals within a population of each species separate foraging activity time along the day or nighttime axis.

However, it is also possible that being active at different times reduces direct, aggressive interactions between individuals of the same species: in a previous study, we found that only rarely did two individuals forage concurrently in a foraging patch and in the infrequent cases when this occurred, 90% of the individuals left within 2 seconds (19).

The question whether the deviation of the internal clock period from 24 hours has any specific function or causal meaning has attracted attention of scientists for years. Traditionally, scientists considered the deviation of tau from 24 hours a simple biological imperfection which is corrected daily by a Zeitgeber (light exposure), and therefore does not lead to serious functional deficits. Accordingly, most studies of this variation focused on its proximate casual mechanism, trying to identify the molecular mechanisms generating the circadian phenotypic diversity, and not on its functional, adaptive significance. However, since the deviation of tau from 24 hours defines the chronotype, one could expect it to be under selection pressure (11). In 1981, Pittendrigh wrote that the deviation is “a strategy, and not a tolerated approximation”, arguing that it contributes to the stabilization of the 24 h rhythms (26). Daan and Beersma (2002), modeled the contribution of the deviation to the stability of the rhythm and concluded that there is no strong theoretical basis for this argument. However, the occurrence of clines in tau along latitudinal gradients suggest the role of natural selection (27, 28). We suggest that this variation reduces intraspecific competition, whether consumptive or aggressive and hence may well be selected for. In fact, variation in tau may reflect a tradeoff between individual dominance in aggressive encounters and foraging efficiency.

Intraspecific morphological variation in the trophic apparatus of different animals has been interpreted as a means for reducing intraspecific resource competition (23, 29). The niche variation hypothesis, first coined by Van Valen (30), suggests that when interspecific selection is relaxed, intraspecific variation increases to reduce intraspecific competition (23). We suggest that the variation in tau that appears in these two species likewise reflects the selective role of intraspecific competition.

In the current study *A. cahirinus* very seldom used torpor, while *A. russatus* entered torpor frequently, and the time spent torpid was correlated not only with individual order of arrival (19), but also with tau. This means that there was a genetic component in the proximate factors that influenced individual chronotype, and that, in turn, determined the ability of the mice to acquire energy and affected their use of torpor as means of reducing energy expenditure. Since previous research suggests that use of torpor in these species results in reduced reproductive activity (22), under the current experimental conditions, where food is supplemented at the beginning of the day and the beginning of the night, earlier chronotypes are expected to have a clear selective advantage. Indeed, previous researchers have suggested that intraspecific temporal partitioning can allow dominant individuals to forage at times of higher rewards, as documented in bellbirds (31) and brown trout (18).

However, under natural conditions with no food supplemented, if the selective advantage depends on food availability, optimal foraging times may vary during the day. Since both species consume mainly arthropods (especially during summer) whose availability changes along the day (25), and since environmental conditions influencing energy expenditure vary along the day and between days and seasons, it is possible that tradeoffs occur between the different selective pressures and that different chronotypes will have a selective advantage during different times.

The source of variation in chronotypes of spiny mice is currently unknown. In humans, genetic influences (heritability) appear to account for a significant proportion. For example, 56% of the variability in food timing, particularly breakfast (32), and 68% of sleep timing on free days (33). An even higher heritability was found in great tits (*Parus major*), with h^2^ of 0.86 ± 0.24 (14). Additionally, chronotypes vary with age and sex: in humans chronotype delays from the age of 10 to 20 years (adolescence), and then advances with age, and on average, females have later chronotypes than males until the average age of menopause (34). A similar change in chronotype with age was also documented in Rhesus Monkeys (*Macaca mulatta*) (35), and an opposite trend, with older individuals having earlier chronotypes was found in in rodents (e.g.,(36)). Spiny mice may live for 2-3 years in natural conditions, so it is entirely possible that their tau changes with age.

We know not the ages of the individual spiny mice studied, nor the age structure of the population. While it would be interesting to speculate that the significant intrapopulation variation in tau reflects both selective pressures at the individual level and age-driven individual variation, further research is required to address this question.

In humans, extreme chronotypes are usually interpreted as syndromes (e.g., delayed and advanced sleep syndrome), although some studies suggest that variability in chronotypes may have adaptive significance under natural conditions, which became maladaptive in the modern world. For example, in the Hadza tribe of hunter-gatherers of Tanzania, a variability in chronotype resulted in one or more individuals awake during 99.8% of the time. Having at least one person awake, will reduce the hazards of sleep to the whole group. When group size drops, active sentinel behavior is adopted (37). Thus intraspecific variation in tau may have evolutionary significance for anti-predator vigilance and for coexistence between competing individuals. Possibly, it has other roles that should be further explored.

In sum, our study suggests that the significant variation in tau that is frequently found among individuals is not a mere ‘biological imperfection’ but has evolutionary significance and is selected for; our study system demonstrates its role in reducing intraspecific competition between individuals within a population.

## Methods

### Study design

The research took place on July-October 2016, under semi natural conditions at four 1000m^2^ field enclosures located in the Dead Sea (for details see (19)). We captured 19 Common spiny mice and 22 Golden spiny mice in the area around the enclosures, using Sherman live traps. Based on previous studies in from our laboratory in these enclosures, and according to the spiny mice population density in nature, and specifically in the Ein Gedi area (19, 22, 38, 39), each enclosure was populated randomly with 8-11 individuals, 4-6 individuals of each species, with a sex ratio that is close to 1. The four enclosures, and the two species served as biological replicates (2 species, 4 enclosures, 4-6 individuals of each species in each enclosure: total of 8 populations). We did not have technical replications since there was no experimental manipulation on the spiny mice populations in the enclosures.

When populating the enclosures, each spiny mouse captured was populated in the enclosure that had the least individuals from the captured individual species and sex, until all enclosures were populated. We had no information regarding period length (which was measured only at the end of the experiment) when the enclosures were populated, thus in this regard the individuals were populated completely randomly in the four enclosures.

Mice were implanted with PIT tags (Destron- Fearing, South St. Paul, MN) subcutaneously and iButtons (DS1922L-F5 thermochron; San Jose, CA, USA) in the abdominal cavity. Body and ambient temperatures were measured continuously (every 20 minutes) over four months, using iButtons. The ambient temperature was measured by placing two iButtons under the boulders of each terrace in each enclosure. Foraging activity and order of arrival to auto-monitored food patches (see below) were measured for 14 days in each month under half-moon conditions (7 consecutive days twice a month). The order of entry of each individual was measured for 14 days a month, for four months (43 days in total), and torpor was measured each day, for 100 days in total (considered as repeated measures for each individual, as explained in the Statistical analysis, p.13).

### Monitoring foraging time

We used auto-monitored foraging patches, comprising a plastic tray (25 cm diameter), in which 2L of local soil were mixed with 3 g of broken sunflower seeds (3g seeds per individual). In each enclosure, we placed two round antennas (20 cm diameter) beneath two foraging trays and each antenna was connected to a transceiver (2001F-ISO; Biomark Ltd, Boise, ID), which logged the tag ID and time (in seconds) of each mouse that entered the patch. Solar panels and a generator provided power supply to the transceivers and antennas. The foraging trays were replenished at dawn and at dusk, the soil was sieved and all the remaining seeds were removed.

### Monitoring the Period length (*τ*) of Spiny mice

Mice were trapped on October 2016 and transferred to the laboratory at Tel Aviv University, where they were individually housed in 38X24X13 cm plastic cages in isolated sound proof rooms, with an ambient temperature of 29°C, under constant dark conditions (DD) with food (standard rodent chew) and water *ad libitum* for 10-14 days. Activity was continuously recorded using infrared detectors (Intrusion detector model MH10; Crow group, Kiriyat-Teufa, Israel) connected to a personal computer. Data were collected by a PC at 6-min intervals using software designed for this purpose (ICPC, Netanya, Israel). The period length of each individual was measured once.

### Data and statistical analysis

#### Foraging sequence

Forging sequence in each foraging patch in each enclosure was determined daily according to the first time an individual arrived to the patch. We calculated the mean order of arrival to the food patches for each individual across days.

#### Use of torpor

torpor temperature threshold was defined according to Willis (40). For each day, we calculated individual total time torpid. Torpor bout duration was determined as time between the beginning of the torpor bout until body temperature returned above the torpor threshold. For each individual, we calculated the mean time spent torpid across days.

#### Determining Period length (*τ*)

Actograms were generated using ClockLab (Actimetrics, Wilmette, IL, USA) and chi-square periodograms were used to calculate the periods of free running period length (*τ*) of each individual.

#### Statistical analysis

For each species, we fitted a statistical model to explore how period length (*τ*) of each individual as a continuous covariate has affected its (1) daily order of entry to the food patches and (2) the amount of time it spent torpid.

All statistical models were fitted using Generalized Linear Mixed Effect Models (GLMM) with a Gamma distribution and the identity of each individual and enclosure as nested random factors. Since our data did not fit the test assumptions of regular linear models (such as normality and homogeneity of variances), we fitted our statistical models using Generalized Linear Mixed Effect Models (GLMM) with a Gamma distribution, which is flexible and does not assume homogeneity. To account for the repeated measures of individuals within different enclosures, we included their identity as nested random factors.

Based on AIC (Akaike Information Criteria) criteria, we used the identity link function for all models, except model 1 for golden spiny mice, where we used the log link function. We also used AIC (Akaike Information Criteria) (41) criteria, to select the important fixed and random effects in each model. For all tests, we used the glmer function (lme4 package (42)) in the R software (R Core Team, 2018).). To improve the stability of the statistical models, we run the models with standardized values of the period length (minutes), and de-standardized the effect sized back to the units of minutes.

For each statistical model, we report the slope of the period length, the significance of the effect of the period length as the change in AIC compared to the model without period length as a dependent covariate, and the p-value of the effect sizes as given by the model. Since we used the log link function in model 1 for the golden spiny mice, we report the estimated change in the order of entry for every unit increase in the period length as *e*^*k*^, where *k* is the estimated slope^63^.

